# Solution structure of *Gaussia* Luciferase with five disulfide bonds and identification of a putative coelenterazine binding cavity by heteronuclear NMR

**DOI:** 10.1101/2020.06.30.176909

**Authors:** Nan Wu, Naohiro Kobayashi, Kengo Tsuda, Satoru Unzai, Tomonori Saotome, Yutaka Kuroda, Toshio Yamazaki

## Abstract

*Gaussia* luciferase (GLuc) is the smallest luciferase (18.2kDa; 168 residues) reported so far and is thus attracting much attention as a reporter protein, but the lack of structural information is hampering further application. Here, we report the first solution structure of a fully active, recombinant GLuc determined by heteronuclear multidimensional NMR. We obtained a natively folded GLuc by bacterial expression and efficient refolding using a solubility tag. Almost perfect assignments of GLuc’s ^1^H, ^13^C and ^15^N backbone signals were obtained. GLuc structure was determined using CYANA, which automatically identified over 2500 NOEs of which > 570 were long-range. GLuc is an all-alpha-helix protein made of nine helices. The region spanning residues 10–18, 36-81, 96-145 and containing eight out of the nine helices was determined with a C_α_-atom RMSD of 1.39 Å± 0.39 Å. The structure of GLuc is novel and unique. Two homologous sequential repeats form two anti-parallel bundles made by 4 helices and tied together by three disulfide bonds. The N-terminal helix 1 is grabbed by these 4 helices. Further, we found a hydrophobic cavity where several residues responsible for bioluminescence were identified in previous mutational studies, and we thus hypothesize that this is a catalytic cavity, where the hydrophobic coelenterazine binds and the bioluminescence reaction takes place.

## Introduction

Luciferase (Luc) is a generic term for bioluminescent enzymes that catalyze the oxidation of a substrate, often termed luciferin [1]. Together with GFP, Luc is widely employed as a reporter protein [2–4]. *Gaussia* Luciferase (GLuc) is a luciferase isolated from the marine *Gaussia princeps* [5], which catalyzes a bright blue light by oxidizing coelenterazine. GLuc is the smallest luciferase reported so far with a molecular mass of 18.2 kDa (excluding the secretion tag). Nonetheless, its bioluminescence intensity is strong (200 fold higher than Firefly Luciferase and *Renilla* Luciferase, the two most widely used luciferase), and it is thus considered as a potential ideal reporter protein [6]. Attempts to improve or redesign GLuc’s bioluminescence characteristics included the lengthening of its half-life luminescence [7–9], and the redshift of its light emission peak at 480 nm [10,11], which is absorbed by tissues during *in vivo* applications [12]. However, structural information at atomic resolution is still not available, making the redesign process tedious.

GLuc contains 10 cysteines, and previous studies demonstrated that the natively folded GLuc contains five disulfide bonds. The presence of 5 disulfide bonds increases the risks of misfolding when GLuc is bacterially produced, resulting in a low yield [13]. In order to overcome this misfolding problem, several methods including fusion with pelB leader sequence [7,14], cell-free systems [15], low-temperature expression [16] were reported, but the yield of natively folded GLuc remained insufficient for high-resolution structural studies. We previously developed a Solubility Enhancement Peptide tag (SEP tag [17–19]). We showed that by attaching a SEP tag containing nine aspartic acids to GLuc’s C-terminus, we could increase the solubility of GLuc, resulting in a spontaneous refolding and the formation of native SS-bonds. Indeed, we obtained nearly 1mg of soluble and functional GLuc from a 200 ml of *E*.*coli* cultured in Luria-Bertani (LB) [11,13,20].

Here, we used the SEP-tag fused GLuc construct to produce a sufficient amount of ^15^N and ^13^C uniformly labeled GLuc for NMR studies. Heteronuclear multidimensional NMR spectroscopy enabled over 99% backbone ^1^H, ^13^C, and ^15^N chemical shifts of GLuc to be assigned. Flexible regions and highly stable regions were identified by ^1^H-^15^N heteronuclear NOE [21] and H/D exchange experiments [22]. The three-dimensional structure calculated by using CYANA (ver 3.98 [23]) were determined with a backbone (C_α_) RMSD of 1.39Å±0.39Å (excluding residues in the flexible regions).

## Results

### Expression and purification of GLuc

The natively folded GLuc possesses ten cysteines that form five disulfide bonds, which can be easily misformed when the protein is expressed in *E*.*coli*, and the cysteines are air-oxidized *in vitro*. Here, we used a SEP-Tag, C9D, which solubilizes the protein during air-oxidization and refolding, thereby increasing the yield of natively folded and active GLuc [20]. The final yield of GLuc after tags cleavage and two times HPLC purification (Fig. S1) was 1.5 mg per liter of M9 minimal medium culture, which was sufficient for NMR analysis. GLuc’s identity was confirmed by MALDI-TOF mass (^15^N labeled GLuc, calculated= 19055.8 Da, experimental=19062.5 Da, Fig. S2). To date, the yield of natively folded active GLuc is almost nil when expressed without the C9D tag [20], and the solubilization tag was thus essential to achieve the present amount of protein, though it was removed once the protein was folded into its native conformation.

### NMR analysis

The ^1^H-^15^N HSQC spectrum exhibited dispersed and sharp peaks (Fig. 1), indicating a stable and well-folded structure. Almost all backbone chemical shifts were visible in the heteronuclear NMR experiments, and over 99% of backbone ^1^H, ^13^C and ^15^N resonances of non-proline residues were unambiguously assigned. C136 was the only un-assigned backbone H-N chemical shifts. The broadened signals around residue C136 suggested that the region encompassing the C136/C148 SS-bond was subjected to structural exchange, as suggested by the ^15^N relaxation dispersion of D138 and L140 (Table 1), and thus the C136 H-N pair was undetected.

**Table 1.** *R*_2_-dispersion experiments on GLuc. Differences of effective ^15^N transverse relaxation rates at CPMG rates of 50 Hz and 1 kHz are listed only for peaks that are resolved and their rate difference > 2 1/s. Larger difference was observed for residues in the C-terminal flexible region (indicated by bold letters) indicative of its structural dynamics. The two-state model analysis of CPMG rates of 50, 100, 150, 200, 250, 300, 400, 500, 600, 800, 1000 Hz showed that the estimated exchange rate was 2500 +/- 800 1/s. Because the residues showing R2 dispersion spread around several blocks, we judged farther residue-specific analysis using single exchange rate is unreliable.

| Residue | R2(50 Hz)-R2(1 kHz)<br>1/s |
| --- | --- |
| V12 | 6.66 |
| S16 | 4.84 |
| T21 | 2.47 |
| D26 | 2.12 |
| G28 | 5.60 |
| L37 | 2.88 |
| A47 | 2.61 |
| S61 | 6.93 |
| M69 | 4.87 |
| K70 | 2.07 |
| G75 | 4.78 |
| T79 | 6.43 |
| T125 | 2.26 |
| D138 | 4.46 |
| L140 | 4.07 |
| **T150** | **4.46** |
| **A152** | **10.85** |

**Fig. 1.**
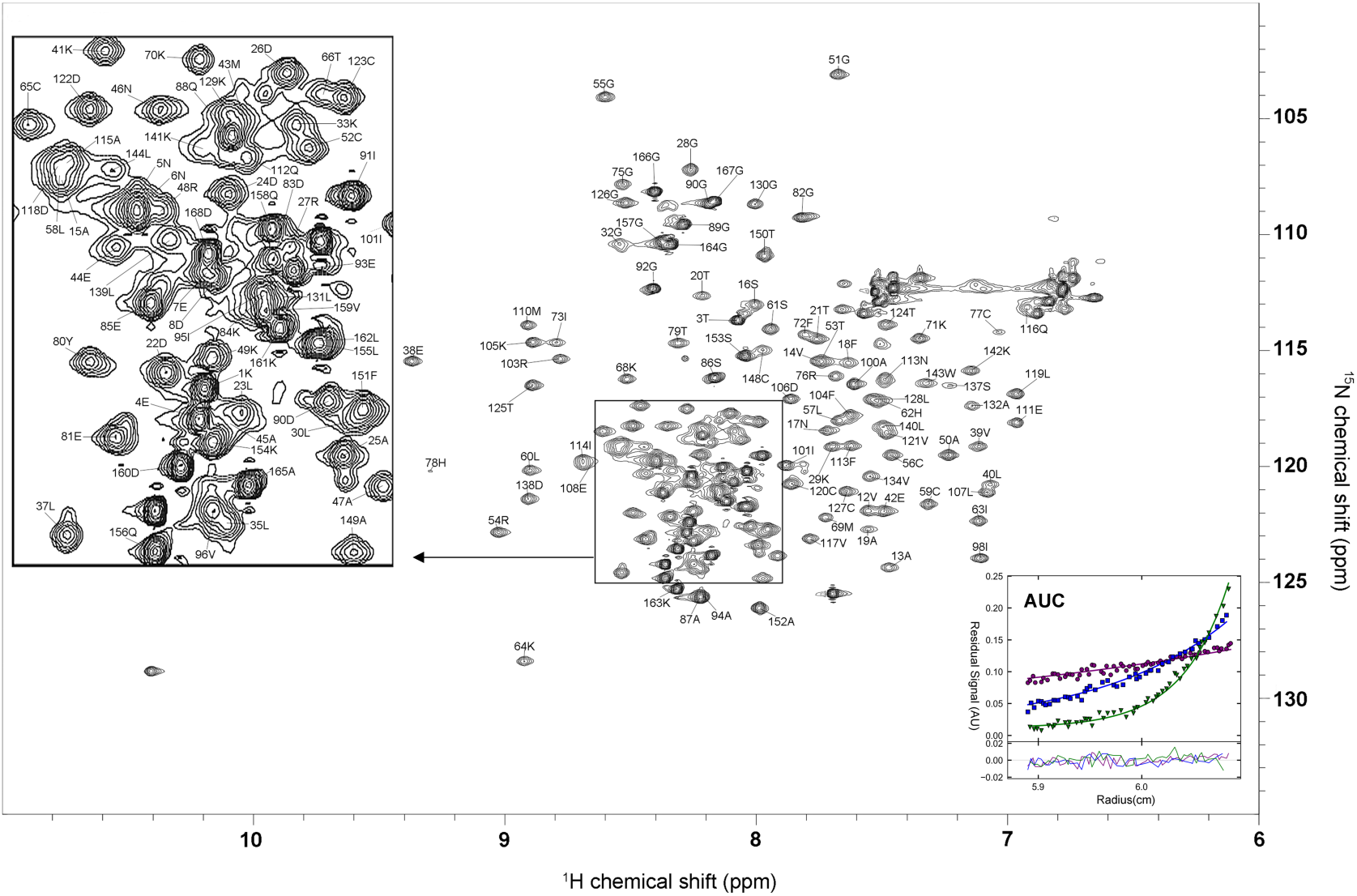
2D ^1^H-^15^N HSQC spectrum of GLuc. The peak assignments are shown using the one-letter code followed by the residue number. Resonance assignments are numbered starting at the first residue (lysine) behind the secretion tag, which was removed without affecting the bioluminescence activity. Mutations at E100A and G103R do not affect activity and are described in our prior paper [13]. The inset in the right bottom (marked AUC) shows the result of sedimentation equilibrium experiments of GLuc protein (concentration at 0.3 mg/mL). Scans from three different rotor speeds (●: 12,000 rpm; ■: 22,000 rpm; ▼: 37,000 rpm) monitored at 280nm. The lines represent the fit to a single species model. The determined molecular weight was 22 kDa, corresponding to a monomer.

The side-chain atoms were automatically assigned by FLYA [24] (a function of CYANA) using the aliphatic atoms identified in the 3D HCCH-TOCSY, ^15^N- and ^13^C-edited NOESY spectra. The assignments were confirmed by visual inspection and when necessary corrected manually using the NMR spectra viewer and analyzer MagRO [25,26]. We assigned over 82.4% of ^1^H, ^13^C and ^15^N atoms of entire GLuc molecule.

The secondary structure elements were analyzed by TALOS+ using the ^1^H, ^13^C, ^15^N chemical shifts (Fig. 2A). TALOS+ indicated that GLuc contains 36.9% helix and 4.7% sheets, in reasonable agreement with our previous prediction based on the consensus of seven publicly available secondary structure predictors (30% helix and 4% sheets) as well as with the results of our Circular Dichroism (CD) analysis (30% helix and 12% sheets) [13]. In addition, the location of helices calculated by TALOS+ and the secondary structure prediction mostly overlapped (Fig. S3).

**Fig. 2.**
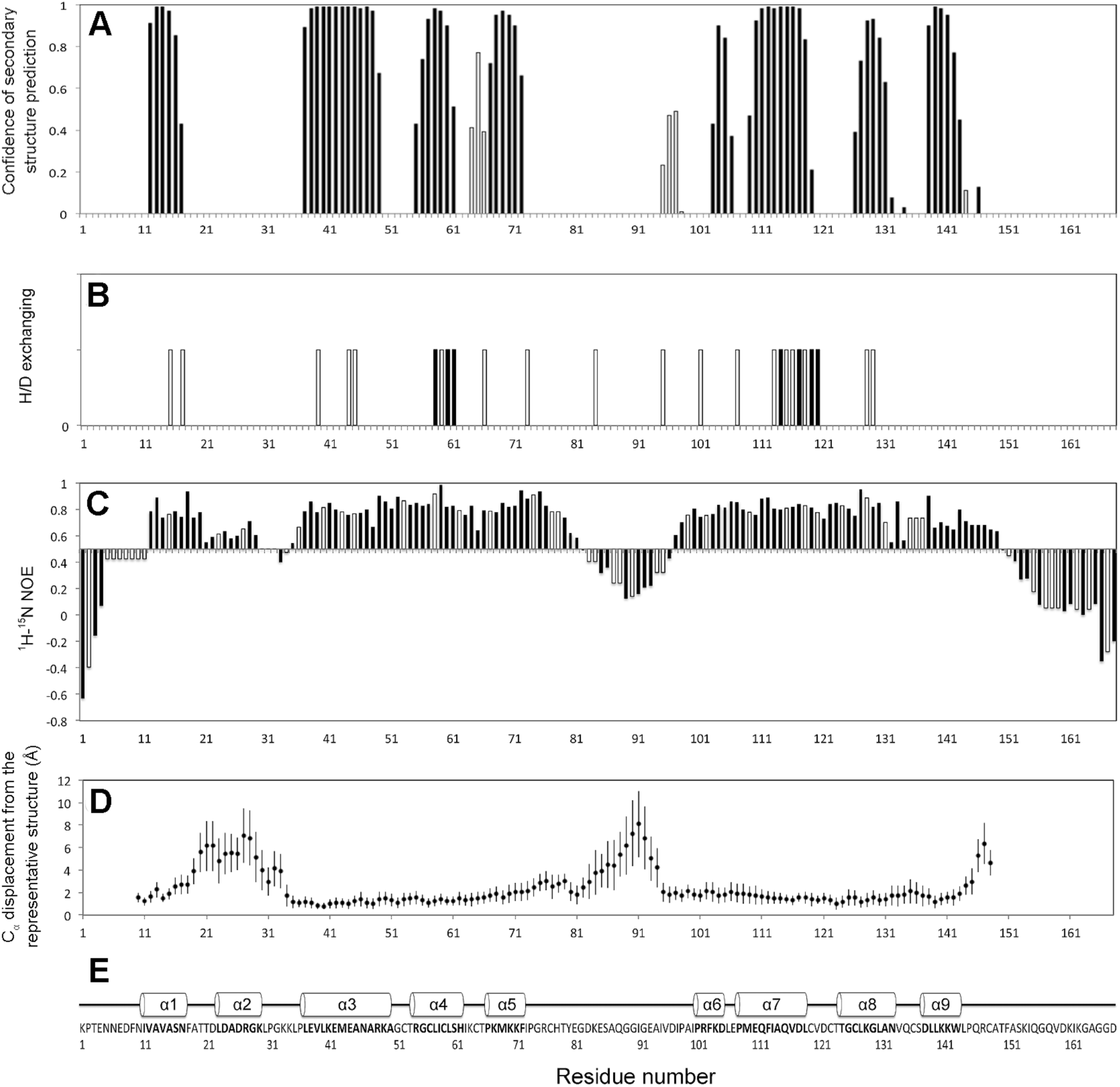
Residue-resolved structural and dynamics features of GLuc. (A) GLuc’s Secondary structure predicted by TALOS+: α-helix and β-sheet tendency are shown with solid and open bars, respectively. (B) H/D exchange experiments. Residues that retained resonance signal after incubation in D_2_O after 20 minutes and 18 hours were marked with open bars and solid bars, respectively (^1^H-^15^N HSQC figures are shown in Fig. S5). (C) ^1^H-^15^N heteronuclear NOE experiment data used to assess GLuc backbone flexibility: ^1^H-^15^N heteronuclear NOE are shown with solid bars. The NOE values of residues that were not identified were assumed using the average value of the preceding and following residues are shown with open bars. Flexible regions of GLuc were identified with the threshold value of 0.5. (D) Backbone C_α_ displacement from the representative structures is calculated using nineteen NMR-derived structures, and the error bars show standard deviations. (E) GLuc’s amino acid sequence and secondary structure that identified from the representative structure.

### Structure calculation and disulfide bond determination

Since GLuc was a monomer as demonstrated by AUC, all NOEs were used as intramolecular NOEs (Fig. 1). The statistics of NOEs assigned during the 19 CYANA runs are shown in Table 2. Even though distance constraints for hydrogen bonds, disulfide bonds, and some manually assigned NOEs were included in the CYANA calculations in addition to the standard automatically assigned NOEs, the target functions were reasonably small (4.07+/-0.59), indicating that the resulting structures were consistent with the experimental data.

**Table 2.** Structural statistics for the nineteen best NMR-derived GLuc structures. 19 rounds of CYANA calculation with different random seed [36]. **\*** The averaged numbers of NOEs and their standard deviations are calculated over the 19 rounds of CYANA calculations. **\*\*** Averaged over the 18 structures (except for the representative structure).

| **NOE distance restraints\*** |  |
| --- | --- |
| All | 2573.4 +/- 42.4 |
| Intra residue ($\vert i-j\vert =0$) | 565.9 +/- 14.7 |
| Sequential ($\vert i-j\vert =1$) | 728.6 +/- 6.6 |
| Medium-range ($2 \leq \vert i-j\vert \leq 4$) | 700.5 +/- 18.1 |
| Long-range ($\vert i-j\vert > 5$) | 579.7 +/- 32.2 |
| Dihedral angle | 183 |
| Hydrogen bonds | 25 |
| Disulfide bonds | 3 |
| **RMSD to the representative structure\*\*** |  |
| Backbone C$_{\alpha}$ (residues 10–18, 36–81, 96–145) | 1.39 Å +/- 0.39 Å |
| Heavy atoms (residues 10–18, 36–81, 96–145) | 1.84 Å +/- 0.20 Å |
| Backbone C$_{\alpha}$ (helices without $\alpha 2$) | 1.30 Å +/- 0.36 Å |
| Heavy atoms (helices without $\alpha 2$) | 1.74 Å +/- 0.45 Å |
| **Ramachandran plot (representative structure)** |  |
| Residues in most favored regions | 71.8 % |
| Residues in additionally allowed regions | 20.4 % |
| Residues in generously allowed regions | 4.9 % |
| Residues in disallowed regions | 2.8 % |

Ellman’s assay indicated that all ten cysteines (C52, C56, C59, C65, C77, C120, C123, C127, C136, and C148) are oxidized in the active GLuc, and thus that they should form five disulfide bonds. Three disulfide bonds C59/C120, C65/C77, and C136/C148 were unambiguously visible in the NMR structures, but the pairing of the remaining four cysteines (C52, C56, C123, and C127), which were close to each other, was less straightforward to determine. In order to determine the remaining two disulfide bonds, we set the distance between the gamma sulfur (Sγ) to > 2Å so that any cysteine could freely combine with any of the remaining three cysteines. We then identified cysteine pairs with Sγ distance < 3Å in the 380 structures obtained from 19 rounds. As a result, C52/C127 and C56/C123 were the most favored pairs and were observed in 92.4% and 56.1% of the calculated structures, respectively. On the other hand, C52/C56 and C123/C127 were observed in only 13.7% and 19.5% of the structures.

### Overall fold and dynamic features of GLuc

The structure with the lowest overall target function among the 20 NMR-derived structures in each round was selected. Nineteen structures from 19 rounds were used for further analysis. Among the nineteen structures, seven structures formed the putatively correct disulfide bonds (C52/C127, C56/C123, C59/C120, C65/C77, C136/C148). We finally selected the structure with the lowest average pairwise RMSD (against all other eighteen structures) as the representative structure. This structure also forms the putatively correct disulfide bonds.

The nineteen superimposed NMR-derived structures with the lowest overall target function show that GLuc has nine helices (α1-α9, Fig. 2E and Fig. 3), and the location of all helices essentially corroborate the TALOS+ prediction except for α2 (Fig. 2A and Fig. 2E). The N-(residues 1-9) and C-terminus (residues 146-168) of GLuc are highly disordered. GLuc’s main structure is formed by residues 10-145, in which the structure of residues10-18, 35-81 and 97-145 were well-defined with an average backbone RMSD to the representative structure for all other eighteen structures of 1.39Å (Table 2). Residues 19-34 and residues 82-96 are highly disordered and can be considered as intrinsically disordered regions (IDR [27], Fig. 2D, and Fig. 3). The structures of helices α1 and α3-α9 were well-defined with an average backbone RMSD of 1.30Å (Table 2). It has been reported that α3-loop-α4-loop-α5 (α3-α5, residues 37-72) and α7-loop-α8-loop-α9 (α7-α9, residues 109-143) are repeat sequences [13,28]. The structure analysis shows that GLuc’s two repeat sequences are connected by the second IDR (residues 82-96) and form an anti-parallel bundle (α3+α8 pair and α4+α7 pair) that surrounds the N-terminal α1 helix. The anti-parallel bundles are firmly tied by three disulfide bonds (C52/C127, C56/C123, and C59/C120), resulting in a high local stability (Fig. 3). Moreover, all residues in the well-defined region exhibited ^1^H-^15^N heteronuclear NOE values larger and more uniform than residues in the N- and C-terminus or in the two IDRs, confirming that the well-defined regions obtained by calculation were consistent with the rigid regions determined by HN NOE values (Fig. 2C and Fig. 2D).

**Fig. 3.**
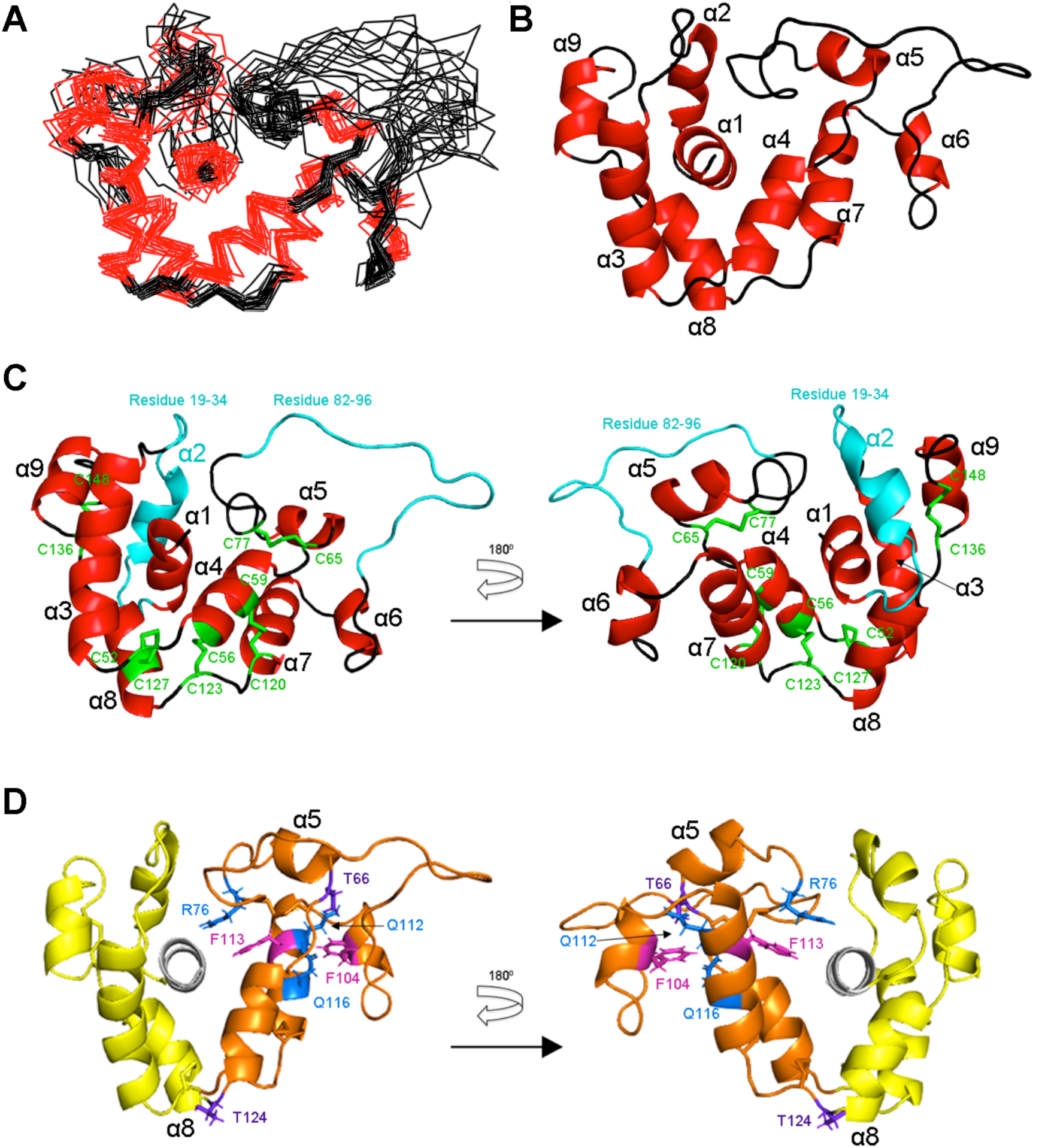
Overall fold of GLuc (residues 10-148) determined by NMR. (A) Wire model of nineteen superimposed NMR-derived structures with the lowest target function. Helices were shown in red and loops in black. (B) Ribbon model of the representative structure. The nine helices of GLuc are marked from α1 to α9. (C) Ribbon model of the representative structure with the five disulfide bonds colored in green. Two IDRs are in cyan. (D) Ribbon model of the representative structure with its two moieties. The tightly packed moiety (residues 52-123) is shown in orange, whereas the loosely packed moiety (residues 19-51, 124-151) is shown in yellow. The central helix, α1 (residues 1-18), is in white. Residues R76, Q112 and Q116 are in blue, F104 and F113 are in magenta, and T66 and T124 are in purple.

## Discussion

The structure of GLuc is novel, as we detected no similar structures in the Protein Data Bank using DALI [29] (Fig. S4). It is even quite different from the structures of *Renilla* luciferase (RLuc) [30], *Oplophorus* Luciferase (OLuc) [31] and apoaequorin [32], which like GLuc uses coelenterazine as a substrate and are ATP independent luciferases. The anti-parallel bundle of helices, which exhibits pseudo 2-fold symmetry, in the GLuc fold can be divided into two moieties. Though both showed well-defined backbone structures, the experimental data indicated differences in the side chain packing stability. The side chains of residues 52-123 are tightly packed whereas those of residues19-51 and 124-151 are loosely packed (Fig. 3D). The high stability of the former one reveals good agreement with the residues showing extreme low H/D exchange rates, whereas residues in the latter one exhibited high H/D exchange rate indicative of a low stability (Fig. 2B and Fig. S5). In the tightly packed moiety, we found several hydrophilic residues with well-determined side-chain structures. For instance, the chemical shifts of R76-Hε, Q112-Hε 1/2, and Q116-Hε 1/2 were clearly different from averaged values observed in a flexible side chain. Furthermore, many NOEs were assigned to these atoms corroborating the fact that these side chains are involved in hydrogen bonds stabilizing the tightly packed moiety. Interestingly the side-chains of R76 and Q112 are stacked to the aromatic rings of F113 and F104, respectively, apparently shifting NMR signals of these protons from ring current effect (Fig. 3D). Finally, the hydroxyl protons of T66 and T124 were also clearly visible, suggesting that they are involved in hydrogen bonds and thus in the N-terminal capping of helices α5 and α8, respectively (Fig. 3D).

Surface accessible analysis of the representative structure indicated a noticeable cavity located among the central α1, α4 and α7 (Fig. 4A, Fig. 4B and Fig. S6). The cavity was made by 19 residues: N10, <u>V12</u>, <u>A13</u>, <u>V14</u>, S16, N17, <u>F18</u>, <u>L60</u>, S61, <u>I63</u>, K64, C65, R76, C77, H78, T79, <u>F113</u>, <u>I114</u>, <u>V117</u> (For reader’s convenience, we underlined the hydrophobic residues; Fig. 4C). Similar cavities formed by these 19 residues were identified in all other eighteen NMR-derived structures, though the sizes and shapes of their cavities showed some variation because of the limited resolution of the NMR structures. The 19 residues are distributed on three structural segments: α4+α7 most rigid block, R76-T79 short loop, and the central α1. The α4+α7 most rigid block was stabilized by the three disulfide bonds as mentioned above (C52/C127, C56/C123, and C59/C120, Fig. 3C). In addition to the rigid structure of the cavity, we observed a structural exchange suggested by ^15^N relaxation dispersion (Table 1). S61 was close to both V12 and T79 (Fig. 4C), which exhibited the second-largest dispersions. We hypothesized that the structural exchange is related to the opened and closed form of this cavity.

**Fig. 4.**
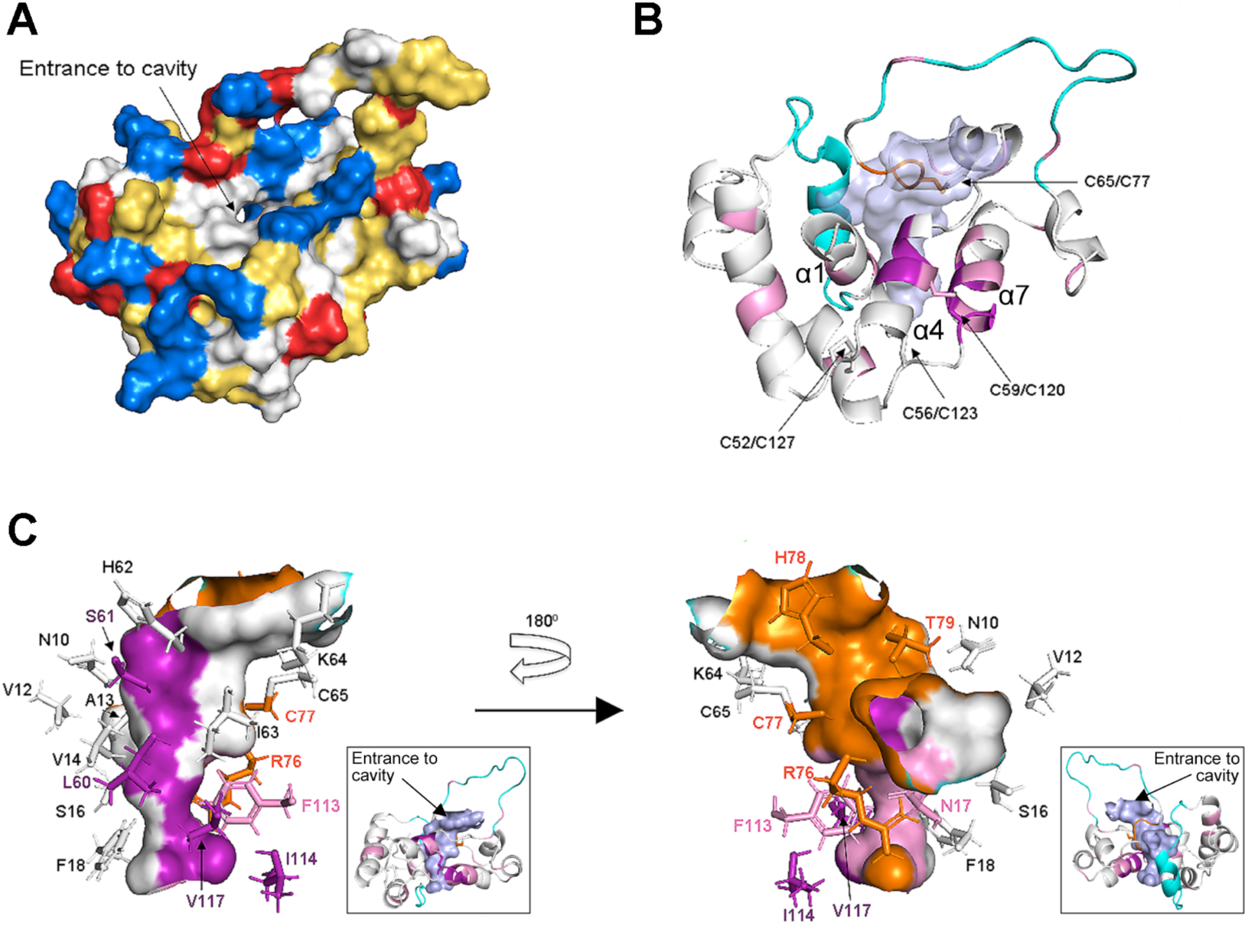
The cavity shown using the representative structure (residues 10-148). (A) Surface representation of GLuc: positive residues (Arg, Lys and His) are colored in blue; negative residues (Glu and Asp) are colored in red; and hydrophobic residues are colored in yellow. The entrance to cavity is indicated by an arrow. (B) The cavity representation (colored in transparent light blue) of GLuc shown from the same direction as in (A). Residues that retained ^1^H-^15^N HSQC signals after 20 minutes and 18 hours H/D exchanging are shown in pink and purple, respectively; residues located in the activity-related loop R76-T79 are in orange; two IDRs are shown in cyan. (C) Residue composition around the interior cavity. The cavity wall and its contributing residues are colored using the same color code as in (B). The insets show ribbon models of GLuc with the cavity colored in light blue and viewed from the same direction as in the main panel. The entrance to the cavity is indicated by an arrow.

A flexible docking simulation indicated that the cavity was large enough to accommodate coelenterazine; and this was verified for all seven models that formed five disulfide bonds (Fig. S7). H/D exchange indicated that the amide protons of N17, L60, S61, F113, I114 and V117, which are located around the cavity, were visible after 20min incubation in D_2_O, and among them, L60, S61, I114, and V117 signals were visible even after 18hrs (Fig. 2B and Fig. S5). The four residues are located in the α4+α7 most rigid block (Fig. 2B and 2E), indicating that the cavity wall is rigid. The hydrophobic character of the cavity’s interior suggests a putative role in recruiting coelenterazine, a small poorly soluble molecule. Furthermore, three activity-related residues R76, C77, H78 [10] are located in the short R76-T79 loop that is near the α4+α7 most rigid block and stabilized through the C65/C77 disulfide bond. C65 and C77 are also associated with bioluminescence activity. In particular, the mutation of C77 resulted in a vanishing luminescence [10]. This can be rationalized by hypothesizing that the destruction of C65/C77 disulfide bond ruins the entire cavity structure and inactivate GLuc.

Sequence alignment also points to the role of the cavity as a binding pocket for coelentarzine. First, the seven cavity forming residues: L60, S61, V117 (in the hydrophobic region); R76, H78 (in the activity-related loop); and C65, C77 (the disulfide bond, see Fig. 4C) are highly conserved in 12 luciferases (MoLuc, MpLuc, etc. see Fig. S8A). C65, R76, C77, V117 were fully conserved, and L60, S61, H78 had a 92% conservation ratio. Furthermore, the structures of OLuc, RLuc and apoaequorin also contain a similar hydrophobic cavity. Altogether, these observations strongly suggest that the cavity constitutes the coelenterazine’s binding site and is thus essential for GLuc’s bioluminescence activity.

Additionally, we noticed that several residues in the C-terminal region (K141-D168) are remarkably conserved (Fig. S8A) despite their high flexibility as assessed by heteronuclear NOE analysis (Fig. 2), which may suggest that they are functionally or perhaps structurally important. The sequence alignment of residues 27-97 with residues 98-168 indicates that K141-F151 in the C-terminal region has a high similarity with K70-Y80 where the aforementioned activity-related loop (R76-T79) is located (Fig. S8B). Furthermore, our previous mutational analysis demonstrated that W143, L144 and F151 also play an important role in GLuc’s activity [11], and it is of interest to note that these conserved residues are disordered in our NMR structure and can be defined as IDRs.

Finally, let us note that several lines of evidence suggested that these residues are not completely disordered. First, residues around F151 exhibited low ^1^H -^15^N NOE values (0.0 ∼ -0.2, Fig. 2C), suggesting a disordered state in the nano- or pico-second time scale, and the *R*_2_-dispersion experiments indicated that these residues experience a micro- or milli-second time scale exchange between the folded and unfolded states rather than in a perfectly flexible state (Table 1). This exchange between a folded and a less folded state was further corroborated by the observation of strong intra-residues and sequential NOEs in the 3D ^15^N-edited NOESY, and the relatively broad line shapes of the peaks in the 2D ^1^H-^15^N HSQC (data not shown). Taken together, our results raise the possibility of an active participation of flexible regions (that can be considered as IDRs) in the coelenterazine oxidation reaction.

## Conclusion

We produced a recombinant ^13^C, ^15^N labeled GLuc in *E*.*coli*, and assigned nearly all of the backbone and most of the side chain chemical shifts. The N- and C-termini, as well as the segment located between α1 and α3 (encompassing α2) and the loop between α5 and α6 were flexible. GLuc’s structure is unique and is made of nine helices, constituting two anti-parallel bundles, which are formed by, respectively, helices α3-α4 and α7-α8 of parts of homologous sequential repeats. The helices are tied together by disulfide bonds to form a 4-finger structure with a pseudo-2-fold symmetry surrounding the N-terminal helix 1. Finally, we identified a hydrophobic cavity where coelenterazine is most likely to bind and the catalytic reaction occurs. The fold of GLuc is novel, and we believe that the above reported structural/dynamic information will open an avenue for redesigning the bioluminescence activity of GLuc and thereby widen its scope of application.

## Materials and methods

### Expression system

A DNA sequence encoding the wild-type GLuc gene (UniProtKB ID: Q9BLZ2) without the 17 residues secretion tag and with an E100A and G103R mutations that increased protein expression was synthesized as reported previously [13]. The GLuc sequence was flanked with an N terminal His-tag and a C terminal SEP-tag (Solubility Enhancement Peptide tag, C9D) to facilitate protein expression, refolding, and purification [20]. Two Factor Xa cleavage sites were inserted between GLuc and His-tag/SEP-Tag. The GLuc gene named GLuc-TG [13] was inserted into pET21c (Novagen) at the NdeI/BamHI site to construct p21GLucTG with ampicillin resistance.

### Protein expression and purification

p21GLucTG was transformed into BL21(DE3), and pre-cultured in 1 L Luria-Bertani (LB) medium at 37°C and 250 rpm shaking. When OD_590nm_ reached 1.0, *E*.*coli* cells were collected by soft centrifugation and transferred to a 1 L M9 medium containing ^13^C-glucose and ^15^NH_4_Cl. Isopropyl β-D-Thiogalactoside (IPTG) was added at 1 mM final concentration for inducing protein expression, and the temperature was lowered to 25°C for minimizing the formation of inclusion bodies. After 4 hours with shaking at 250 rpm, the cells were harvested by centrifugation and sonicated. GLuc was purified from the supernatant fraction using a Nickel Nitrilotriacetic Acid (NTA) column followed with overnight dialysis at 4°C against 50 mM Tris-HCl, pH 8.0. GLuc was then air-oxidized for three days at the same conditions in order to form the five disulfide bridges. Residual misfolded GLuc was removed using a reversed phase High-Performance Liquid Chromatography (HPLC). The protein concentration was determined using a Bradford assay [33], and Factor Xa was added to GLuc dissolved in 50 mM Tris-HCl, 100 mM NaCl, and 5 mM CaCl_2_ at a ratio of 1:100 (w/w), and the sample was again incubated for 8 hours at 37°C, 100 rpm for enzymatic cleavage of the His- and the SEP-Tags. Uncleaved GLuc was removed using, again, reversed phase HPLC. GLuc identity was confirmed by MALDI-TOF mass spectroscopy on an ABI SCIEX TOF/TOF 5800 (Thermo Fisher Scientific Inc., Massachusetts, USA). GLuc was freeze-dried and kept as a powder at -30°C until use.

### Analytical Ultracentrifugation (AUC)

Sedimentation equilibrium experiments were carried out using an Optima XL-A analytical ultracentrifuge (Beckman-Coulter, Inc., Brea, California, USA) with a four-hole An60Ti rotor at 20°C. Before centrifugation, GLuc samples were dialyzed overnight against 50 mM MES and 100 mM NaCl at pH 4.7. The solvent density (1.006983 g/cm^3^) was determined using DMA 5000 (Anton Paar). Each sample was then transferred into a cell with a six-channel centerpiece. The sample concentrations were 1.2, 0.6, and 0.3 mg/mL. Data were obtained at 12,000, 22,000, and 37,000 rpm. A total equilibration time of 24 hours was used for each speed, with absorbance scans at 280 nm taken every 4 hours to ensure that equilibrium had been reached. Data analysis was performed by global analysis of all of the data sets obtained at different concentrations and rotor speeds using SEDPHAT [34].

### NMR analysis and structure calculation

NMR experiments for resonance assignments and ^1^H-^15^N heteronuclear NOE experiment were conducted using 0.2 mM ^15^N single or ^15^N, ^13^C double labeled GLuc protein dissolved in 50 mM MES buffer pH 6.0 and 2 mM NaN_3,_ at 293 K with 8%(v/v) D_2_O in a 5 mm Shigemi microtube (Shigemi co., Ltd, Tokyo, Japan). NMR spectra were acquired on a Bruker Avance-III 700 MHz spectrometer, equipped with a 5 mm CPTXI cryoprobe. Two-dimensional and three-dimensional NMR experiments (^1^H-^15^N HSQC, HNCACB, CBCA(CO)NH, HNCA, HNCO, HN(CA)CO) were performed for the backbone ^15^N and ^13^C assignments. ^15^N-TOCSY-HSQC, ^15^N-NOESY-HSQC, and HCCH-TOCSY were used for backbone and side-chain signal assignments. H/D exchange ^1^H-^15^N-HSQC experiment was performed under the above-described conditions but by dissolving GLuc’s freeze-dried powder in D_2_O instead of H_2_O. The transverse relaxation rate (*R*_2_) dispersion experiments for backbone ^15^N atoms were performed on a Bruker Avance-III 900 MHz spectrometer, using pulse scheme including constant time relaxation compensated CPMG pulse sequences [35]. A series of 2D experiments were acquired with various rates of CPMG pulse (50, 100, 150, 200, 250, 300, 400, 500, 600, 800, 1000 Hz) in the two of 20 ms CPMG blocks. The ^15^N CPMG irradiation intensity was set to 3125 Hz (∼35 ppm). To avoid the error in peak intensity arisen from the offset effect, two sets of relaxation experiments with different ^15^N irradiation centers (111 ppm and 125 ppm) were carried out. For each signal, we read the series of peak intensities using the spectrum giving the smaller errors by the offset effect.

### Three-dimensional structure determination

Automated NOE assignments and structure calculations were performed using CYANA (ver. 3.98) on a PC-cluster equipped with 20-core Intel Xeon E5-4627v3 (3.0 GHz) and using the manually assigned chemical shifts and a list of NOE chemical shifts derived from the 3D ^15^N-, ^13^C-edited NOESY spectra of the aliphatic and aromatic regions. For each cycle of CYANA calculation, 20 out of 100 structures were selected after 10,000 steps of simulated annealing using distance constraints derived from automatically assigned NOEs. The detailed algorithm and strategy are described [23]. For the automated NOE assignments of the NOEs peaks, the tolerances were set to 0.04, 0.4 and 0.4 ppm for ^1^H, ^15^N and ^13^C signals, respectively. 19 rounds of CYANA calculations with different random seeds were performed. It should be noted that, for a minor number of the NOEs, the automated assignment was ambiguous and depended on the random seed. We thus calculated the structures using restraints that were slightly different from set to set, and we selected the best structure from the structures generated using each of the sets and reported the ensemble of structures. The treatment using many random seeds was previously discussed and the ensemble of structures calculated using ambiguous assignments becomes more diverse and safer. We can reduce the risk to make structure ensemble affected by wrong NOE assignments [36]. According to experimentally determined slow-exchanging backbone amide protons and threonine (Thr) side chain OH atoms, we applied 24 sets of distance constraints, two of them for hydrogen bonds including Thr-OH related to N-terminal capping of α-helices and 21 of them for backbone amide protons to the 19 rounds of CYANA calculations.

### Other softwares

The secondary structure elements of GLuc were predicted by submitting the backbone chemical shifts into TALOS+ (https://spin.niddk.nih.gov/bax/nmrserver/talos/) [37]. Root-Mean-Square Deviation (RMSD) was calculated using a Biopython.PDB module (https://biopython.org) [38]. Three-dimensional images of the NMR structures were generated using PyMOL. The flexible docking simulation was calculated using AutoDock (Ver. 4.2.6) and AutoDockTools (Ver.1.5.6) [39]

## Acknowledgments

We thank members of the Kuroda Laboratory for discussion, and help and advice with the experimental operations. We are especially thankful to Dr. Tetsuya Kamioka and Mr. Fumiya Suzuki for advice and kind help with protein expression and purification. We are grateful to Prof. Russel Hopcroft, University of Alaska–Fairbanks (UAF), for kind permission to reproduce his photograph of *Gaussia princeps* (used in the graphical abstract; R. Hopcroft copyright), and Prof. Yanhong Bai, Zhengzhou University of Light Industry, for her generous support of this international joint research.

## Authors’ contribution

W.N, Y.K. and Y.T. conceived the project, analyzed the structure, and wrote the manuscript. W.N., K.T., and T.Y. performed the NMR experiments and analyzed the data, N.K and T.Y. performed and analyzed the structure calculation with W.N’s assistance, S.U. and T.S. performed the AUC experiments. All authors contributed to finalize the manuscript and approved it.

## Database

The chemical shifts have been deposited in the Biological Magnetic Resonance Bank (BMRB) under the accession <u>No.36288</u>, and the atomic coordinates are deposited in the Protein Data Bank under accession number PDB-ID: <u>6KYN</u>. The expression vector for GLuc-TG (p21GLucTG) is deposited in Addgene (<u>ID:124660</u>).

