## Supplementary material for "Solution structure of *Gaussia* Luciferase with five disulfide bonds and identification of a putative coelenterazine binding cavity by heteronuclear NMR": Fig. S

**Supplementary Figures**

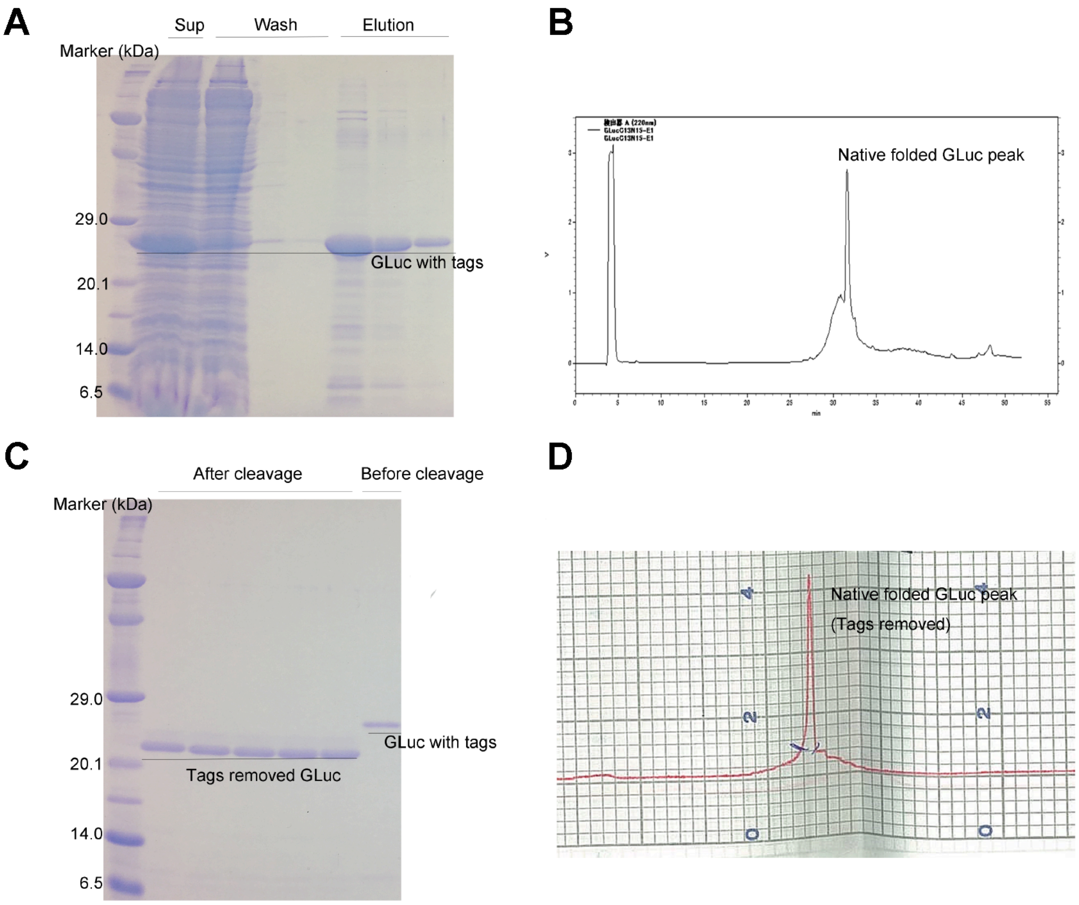


**Fig. S1. Expression and Purification of GLuc (labeled with $^{13}$C and $^{15}$N)** The cultured *E.coli* cells were harvested by centrifugation and the pellet was sonicated. GLuc was purificated by applying the supernatant fraction to a Nickel NTA column followed with washing (three times) and elution (three times). An aliquot (0.05% of 1 liter culture) from the the supernatant (Sup) after sonication, washing and elution was collected for the SDS-PAGE analysis (A). The eluted GLuc was dialyzed and purified by reverse phase HPLC in order to collect the main peak that was the natively folded GLuc fraction (B). The collected peak portion was freeze dryed and then dissolved in MilliQ water for Factor Xa cleavage. An aliquot of the samples before and after (five parallel experiments) cleavage was analyzed by a SDS-PAGE (C). The tags removed GLuc was then purified again through a $2^{nd}$ passage to the reverse phase HPLC (D). The main peak was collected in order to keep all GLuc sample have a uniform native fold. Futher purification details are given in [1].

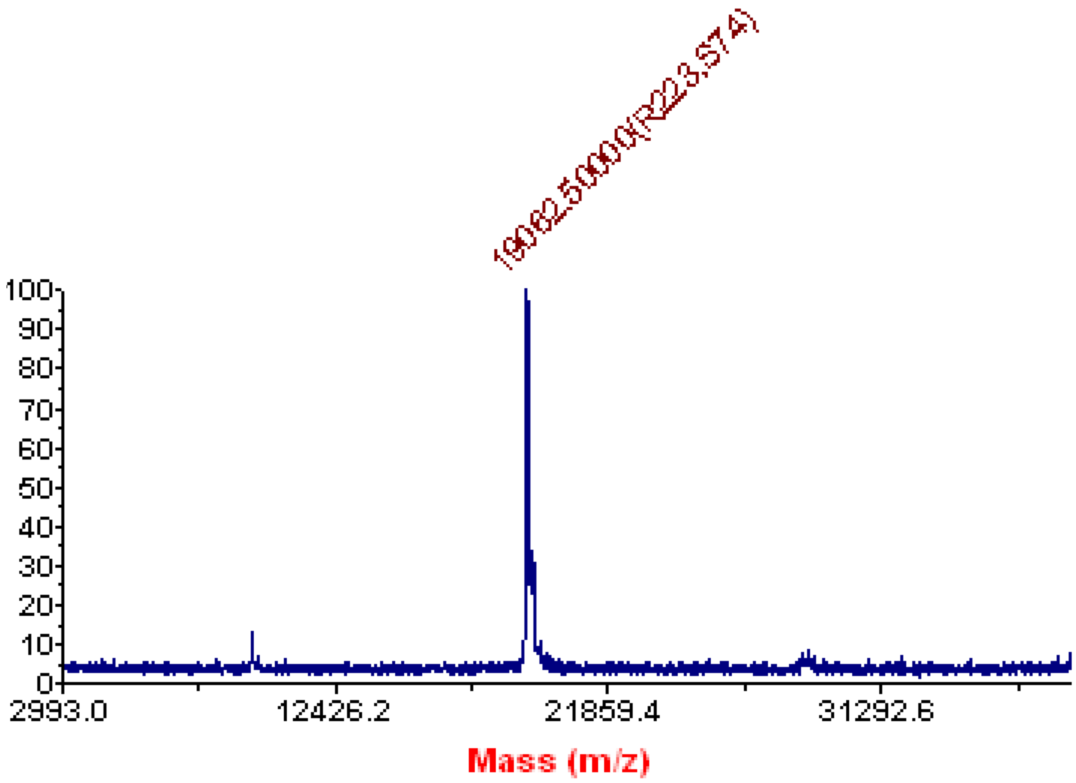


**Fig. S2. MALDI-TOF mass of $^{15}$N labeled GLuc expressed in *E.coli*.** Freeze dryed $^{15}$N labeled GLuc was dissolved in a matrix solution (1mL matrix solution contains: 10 mg sinapic acid, 500 μL acetonitrile, 100 μL 1% TFA and 400 μL MilliQ) at a final protein concentration of 5 μM. One microliter (containnig 5 pmol of GLuc) was loaded on the MALDI-TOF plate for measurement. The calculated mass of $^{15}$N labeled GLuc is 19055.8 Da.

```
                    ....|....|....|....|....|....|....|....|....|....|
                        5    10   15   20   25   30   35   40   45   50
GLuc sequence       KPTENNEDFNIVAVASNFATTDLDADRGKLPGKKLPLEVLKEMEANARKA
TALOS+              cccccccccchhhhhhhcccccccccccccccccchhhhhhhhhhhhhhc
Consensus 85%       cccccccc--eee--ccc-----ccc-cccccccchhhhhhhhhhhhhhc
NOE flexible        ≈≈≈≈≈≈≈≈≈≈≈
H/D 20min                         * *                     *    **
H/D 18h
Packing status                        ○○○○○○○○○○○○○○○○○○○○○○○○○○○○○○○○
Domain(repeats)                               111111111111111111111111

                    ....|....|....|....|....|....|....|....|....|....|
                        55   60   65   70   75   80   85   90   95  100
GLuc sequence       GCTRGCLICLSHIKCTPKMKKFIPGRCHTYEGDKESAQGGIGEAIVDIPE
TALOS+              cecchhhhhhhcceechhhhhhcccccccccccccccccccccceeeccc
Consensus 85%       cccc--eee--ccccc-c----ccccc---ccccc-ccccc---ee-ccc
NOE flexible                                       ≈≈≈≈≈≈≈≈≈≈≈≈≈≈≈
H/D 20min                  ****    *      *          *          *
H/D 18h                    + ++
Packing status      ○●●●●●●●●●●●●●●●●●●●●●●●●●●●●●●●●●●●●●●●●●●●●●●●●●
Domain(repeats)     11111111111111111111111111111111111111111111111222

                    ....|....|....|....|....|....|....|....|....|....|
                       105  110  115  120  125  130  135  140  145  150
GLuc sequence       IPGFKDLEPMEQFIAQVDLCVDCTTGCLKGLANVQCSDLLKKWLPQRCAT
TALOS+              cchhhhcchhhhhhhhhhhhccccchhhhhhhhcccchhhhhhhccccccc
Consensus 85%       cccccccchhhhhhhhhh---hh--hhhh-------hhhhhhh------h
NOE flexible                                                         ≈
H/D 20min           *     *     ********       **
H/D 18h                          +  + ++
Packing status      ●●●●●●●●●●●●●●●●●●●●●●●○○○○○○○○○○○○○○○○○○○○○○○○○○○
Domain(repeats)     22222222222222222222222222222222222222222222222222

                    ....|....|....|....|
                       155  160  165  170
GLuc sequence       FASKIQGQVDKIKGAGGD
TALOS+              cccccccccccccccccc
Consensus 85%       hhhhhh--hh----cccc
NOE flexible        ≈≈≈≈≈≈≈≈≈≈≈≈≈≈≈≈≈≈
H/D 20min
H/D 18h
Packing status      ○
Domain(repeats)     222222222222222222


h = 4-turn helix (α helix). Min length 4 residues.
e = extended strand in parallel and/or anti-parallel β-sheet conformation. Min length 2 residues.
c = coil (residues which are not in any of the above conformations).
```

**Fig. S3. Residue-level structural characterization of GLuc.** TALOS+ shows GLuc's secondary structure calculated from the chemical shifts using the TALOS+ server [2]. Consensus 85% indicates GLuc secondary structure predicted using seven publicly available predictors with 85% consensus [3]. NOE flexible and marked with "≈" shows highly flexible residues identified by $^{1}H$-$^{15}N$ heteronuclear NOE experiment. H/D 20min and H/D 18h respectively shows residues that retained their signal after 20 minutes (marked with "*") and 18 hours (marked with "+") in $D_2O$. Packing status shows the tightly packed moiety (marked with "●") and the loosely packed moiety (marked with "○"), respectively. Domain (repeats) indicates the two homologous repeats of GLuc that identified in the reference [4].

```
MODEL A
# No:  Chain   Z    rmsd lali nres  %id PDB  Description

MODEL B
# No:  Chain   Z    rmsd lali nres  %id PDB  Description
   1:  4ddg-A  2.5  3.5   59   399    5   MOLECULE: UBIQUITIN-CONJUGATING ENZYME E2 D2, UBIQUITIN THI
   2:  2xhi-A  2.4  3.7   52   316   10   MOLECULE: N-GLYCOSYLASE/DNA LYASE;
x  3:  4oge-A  2.4  9.3   99   977    5   MOLECULE: HNH ENDONUCLEASE DOMAIN PROTEIN;
x  4:  5ukh-A  2.3  5.6   52   321   13   MOLECULE: UNCHARACTERIZED PROTEIN;
   5:  1u9p-A  2.3  3.6   48    96    4   MOLECULE: PARC;
   6:  2rji-A  2.3  2.8   53    84    2   MOLECULE: ERYTHROCYTE BINDING ANTIGEN 175;
   7:  4d8o-A  2.3  3.0   46   509   11   MOLECULE: ANKYRIN-2;
   8:  3mzy-A  2.2  4.0   67   123    9   MOLECULE: RNA POLYMERASE SIGMA-H FACTOR;
   9:  5dic-A  2.2  2.9   51   115    4   MOLECULE: ODORANT-BINDING PROTEIN;
  10:  6e11-E  2.2  4.2   65   210    8   MOLECULE: UNKNOWN (CLAW);
x 11:  5gha-D  2.1  5.6   55   310    4   MOLECULE: SULFUR TRANSFERASE TTUA;
x 12:  5bmq-A  2.1  7.4   59   205    8   MOLECULE: ERFK/YBIS/YCFS/YNHG FAMILY PROTEIN;
x 13:  4dbg-B  2.1  8.2   50   146   10   MOLECULE: RANBP-TYPE AND C3HC4-TYPE ZINC FINGER-CONTAINING
  14:  2g7r-A  2.1  2.7   48    86   13   MOLECULE: MUCOSA-ASSOCIATED LYMPHOID TISSUE LYMPHOMA TRANSL
x 15:  6hls-A  2.1 10.5   70  1431   11   MOLECULE: DNA-DIRECTED RNA POLYMERASE I SUBUNIT RPA190;
  16:  3oao-A  2.0  4.2   47   140    9   MOLECULE: UNCHARACTERIZED PROTEIN FROM DUF2059 FAMILY;
               H1 H2 H3 H4 H5 H6 H7 H8 H9
       4ddg-A  o  -  o  -  -  -  -  o  -
       2xhi-A  o  -  o  o  -  -  -  -  -
       1u9p-A  o  -  o  o  -  -  -  -  -
       2rji-A  o  -  o  -  -  -  -  o  -
       4d8o-A  o  -  o  o  -  -  -  -  -
       3mzy-A  o  -  o  o  -  -  -  o  -
       5dic-A  -  -  o  o  -  -  -  o  -
       6e11-E  o  o  o  -  -  -  -  o  -
       2g7r-A  o  -  o  o  o  -  -  -  -
       3oao-A  o  o  o  o  -  -  o  -  -

MODEL C
# No:  Chain   Z    rmsd lali nres  %id PDB  Description
   1:  4qo5-A  2.4  4.8   73   521    8   MOLECULE: HYPOTHETICAL MULTIHEME PROTEIN;
   2:  5c4y-A  2.2  2.7   61   136    3   MOLECULE: PUTATIVE TRANSCRIPTION REGULATOR LMO0852;
   3:  2g7r-A  2.0  2.8   49    86   12   MOLECULE: MUCOSA-ASSOCIATED LYMPHOID TISSUE LYMPHOMA TRANSL
               H1 H2 H3 H4 H5 H6 H7 H8 H9
       4qo5-A  -  o  o  o  o  -  o  -  -
       5c4y-A  -  o  o  o  o  -  o  -  -
       2g7r-A  o  -  o  o  o  -  -  -  -

MODEL D
# No:  Chain   Z    rmsd lali nres  %id PDB  Description

MODEL E
# No:  Chain   Z    rmsd lali nres  %id PDB  Description
   1:  5k78-B  2.8  3.7   61   351   10   MOLECULE: RNA LARIAT DEBRANCHING ENZYME, PUTATIVE;
   2:  2r18-A  2.0  2.3   41   129   10   MOLECULE: CAPSID ASSEMBLY PROTEIN VP3;
               H1 H2 H3 H4 H5 H6 H7 H8 H9
       5k78-B  o  -  o  -  -  -  -  o  -
       2r18-A  -  -  o  -  -  -  -  -  o

MODEL F
# No:  Chain   Z    rmsd lali nres  %id PDB  Description
   1:  4d8o-A  2.1  3.0   47   509    9   MOLECULE: ANKYRIN-2;
   2:  2h56-A  2.0  3.1   45   218   11   MOLECULE: DNA-3-METHYLADENINE GLYCOSIDASE;
               H1 H2 H3 H4 H5 H6 H7 H8 H9
       4d8o-A  o  -  o  -  -  -  -  o  -
       2r18-A  -  -  o  -  -  -  -  -  o

MODEL G
# No:  Chain   Z    rmsd lali nres  %id PDB  Description
x  1:  1e3a-B  2.8  6.5   85   560    8   MOLECULE: PENICILLIN AMIDASE ALPHA SUBUNIT;
```

**Fig. S4. Similar structure search of GLuc against the Protein Data Bank using Dali server.** Seven GLuc models that have all five disulfide bonds were selected from the nineteen NMR-derived structures and submitted to a Dali server (http://ekhidna2.biocenter.helsinki.fi/dali/) [5] to search similar structure in the Protein Data Bank. The data outputs of Dali sever of all seven GLuc models are shown for MODEL A (representative structure), B, C, D, E, F and G, respectively. None the searches showed a similar structure for MODEL A and D. Although a few similar structures were identified for on MODEL B, C, E, F and G, all of which exhibited low Z-score (<3.0, which over 6.0 could be taken to be highly significant), indicating no significant structural similarity. Structures marked with a “×” were firstly ignored, because they have an rmsd >4.0 are structurally dissimilar to GLuc. Structures with rmsd less than 4.0 were then partially matched with GLuc models over helix segments (shown in “Details”, “H1”-“H9” represent nine helices of GLuc, all matched helices were marked with “○” below). GLuc structure shows H3H4H5-H7H8H9 is critical for GLuc’s structural skeleton (Fig. 3) and the putative catalytic cavity (Fig. 4), but none of the identified structures matched GLuc in this areas, which strongly suggested that the matches are accidental.

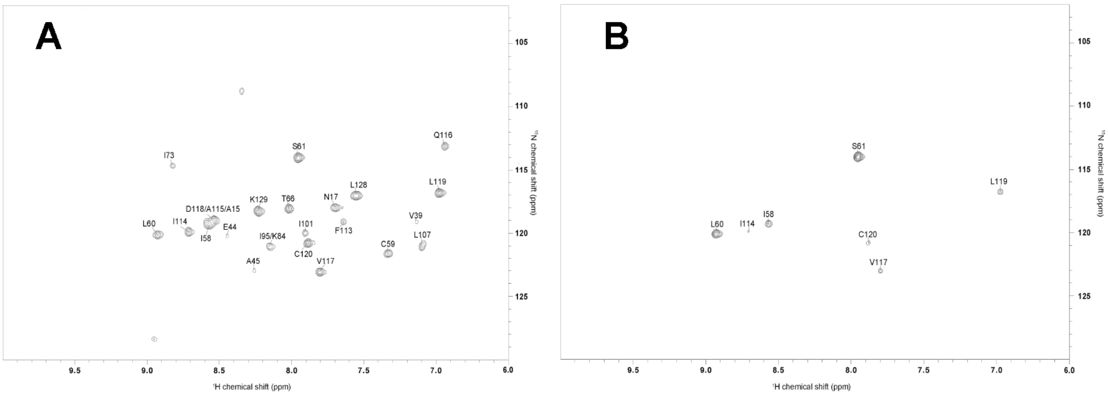


**Fig. S5. H/D exchanging $^{1}H$-$^{15}N$ HSQC spectra of GLuc.** Measurements were initialed by dissolving GLuc to a final concentration of 0.2 mM in $D_2O$ containing 50 mM MES buffer pH 6.0 with 2 mM $NaN_3$. $^{1}H$-$^{15}N$ HSQC spectra were measured after incubating GLuc at 298 K (A) for 20 minutes and 18 hours (B), respectively.

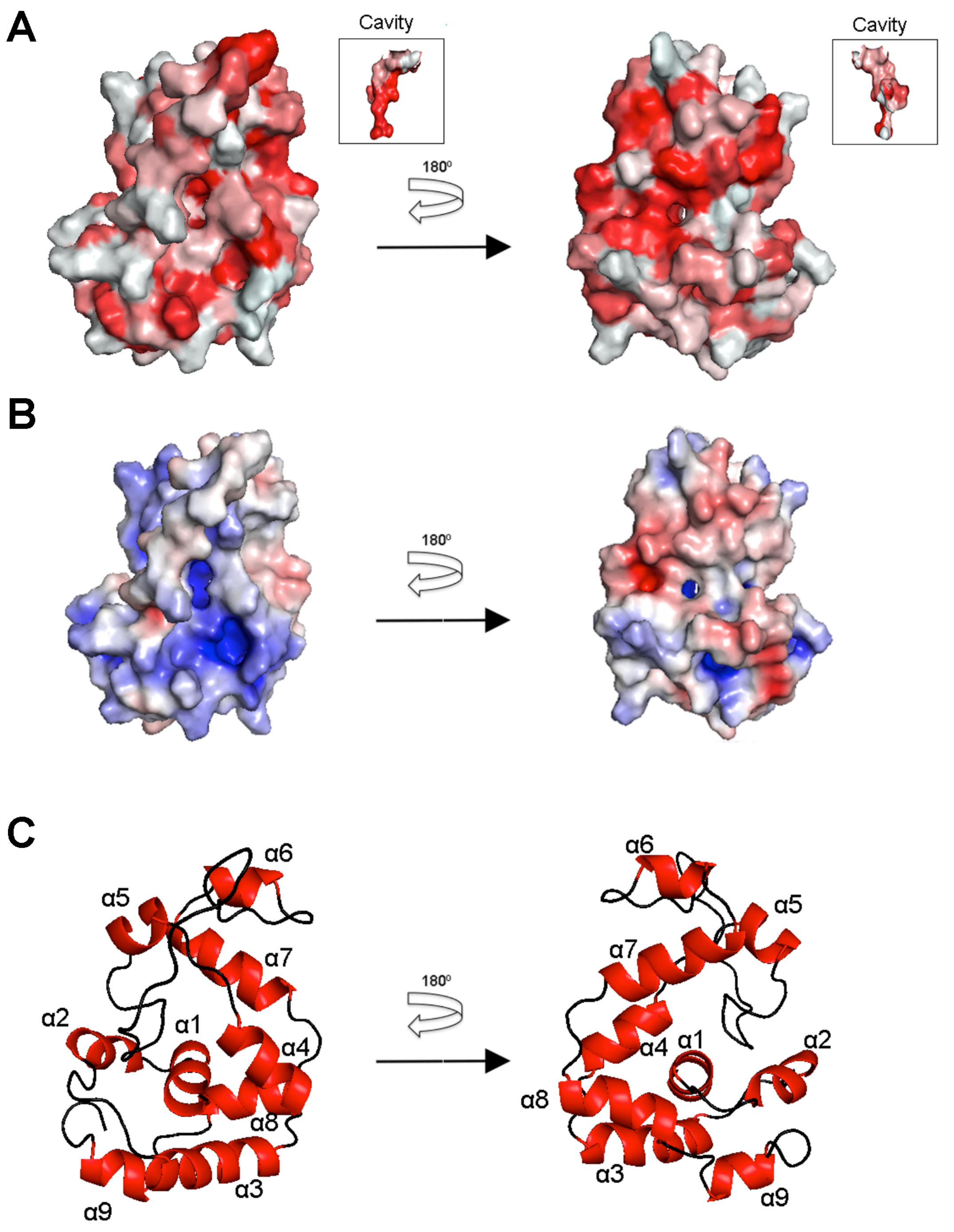


**Fig. S6. Surface representation of GLuc (representative structure, residues 10-148).** (A) Hydropathy surface representation of GLuc shown by coloring the molecule using "color_h" pymol script (https://pymolwiki.org/index.php/Color_h): hydrophilic regions were shown in white and hydrophobic regions were shown in red. The insert figures show the interior cavity. (B) Electrostatic surface representation of GLuc calculated using an APBS plugin of Pymol.

Positive charges are shown in blue, negative charges are shown in red and neutral residues are in gray. (C) Ribbon representation of GLuc from the same direction as in (A) and (B).

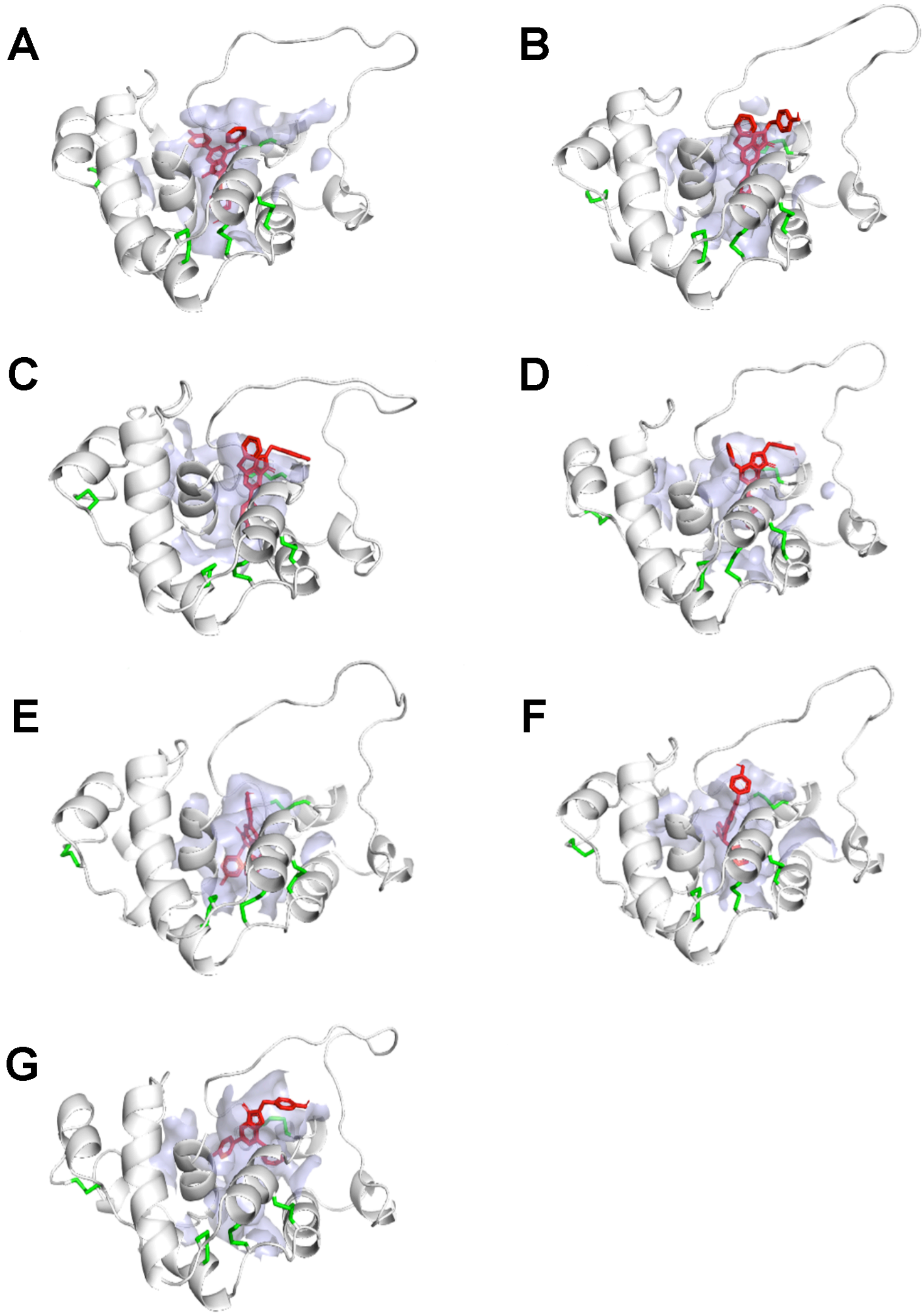


**Fig. S7. Conformations of seven best docking complexes.**

Seven GLuc models that are referred in Fig. S4 were docked with coelenterazine using Autodock4.2 [6]. The coelenterazine and three activity-related residues on GLuc (R76, H78 and T79) were set as flexible, and all other residues were set as rigid. 50 docking complexes were generated during the docking simulation of each GLuc model (Genetic Algorithm), and the one with lowest binding energy were selected and shown in (A) ~ (G), respectively. (A) shows the docking complex of MODEL A (representative structure), binding energy= -6.47 kcal/mol, inhibit constant= 17.96 μM; (B) shows the docking complex of MODEL B, binding energy= -6.13 kcal/mol, inhibit constant= 32.26 μM; (C) shows the docking complex of MODEL C, binding energy= -5.8 kcal/mol, inhibit constant= 56.21 μM; (D) shows the docking complex of MODEL D, binding energy= -7.07 kcal/mol, inhibit constant= 6.57 μM; (E) shows the docking complex of MODEL E, binding energy= -9.29 kcal/mol, inhibit constant= 154.09 nM; (F) shows the docking complex of MODEL F, binding energy= -11.8 kcal/mol, inhibit constant= 2.26 nM; (G) shows the docking complex of MODEL G, binding energy= -8.36 kcal/mol, inhibit constant= 742.04 nM. GLuc molecules are shown in white; coelenterazines are shown in red; five disulfide bonds are shown in green; the interior surface (cavity) that is formed by residues N10, V12, A13, V14, S16, N17, F18, L60, S61, I63, K64, C65, R76, C77, H78, T79, F113, I114, V117 are shown in transparent light blue.

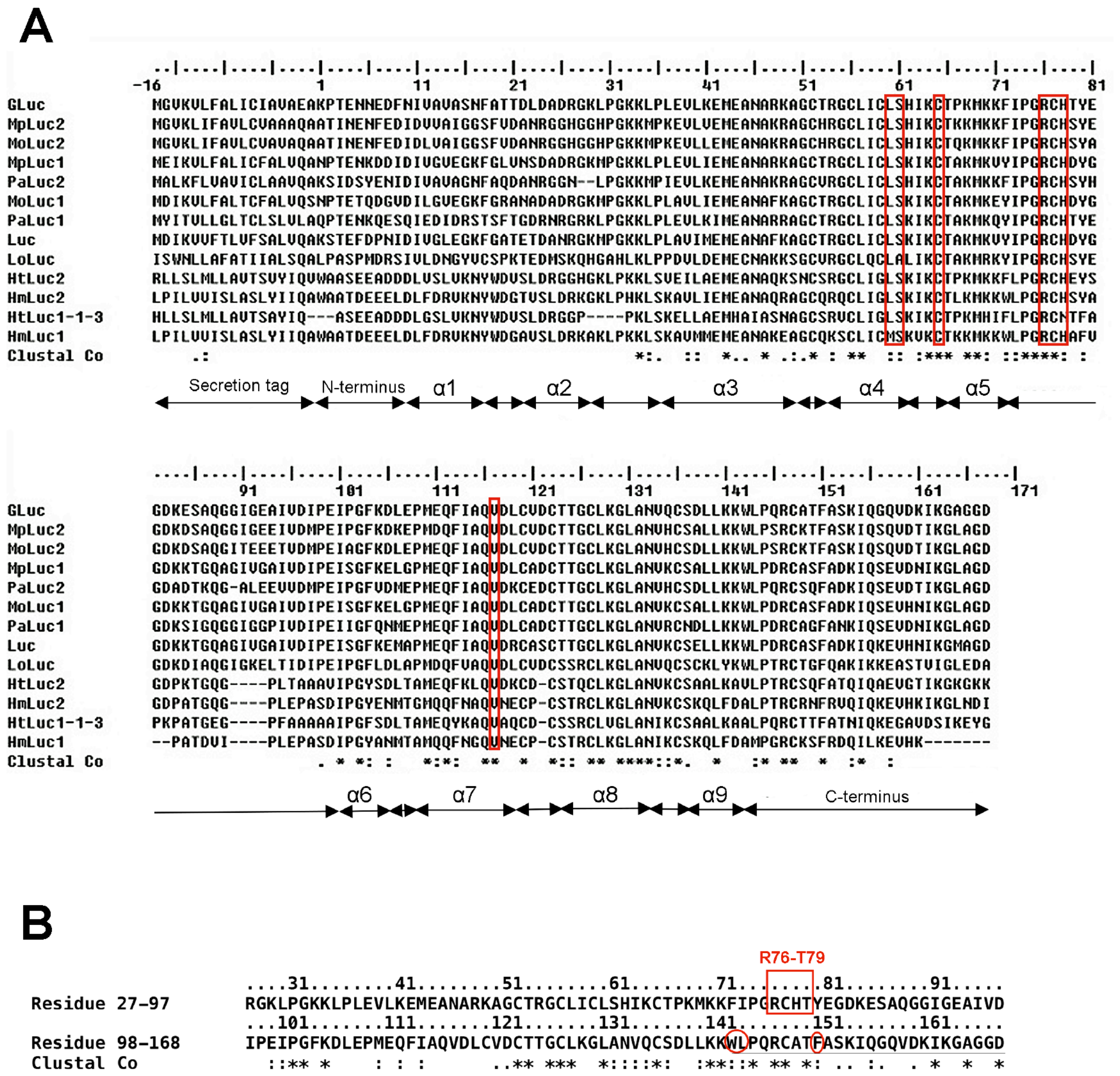


**Fig. S8. Sequence alignment of luciferases.**

GLuc alignment with 12 similar luciferases is shown in (A). GLuc indicates the GLuc's amino acid sequence. Residues -16 to 0 represent a secretion tag sequence. Clustal Co indicates amino acid conservation throughout the 12 sequences of BLAST-detected luciferases according to ClustalW [3]: (An asterisk indicates a fully conserved residue, a column shows a highly conserved residue, and a dot shows a poorly conserved residue according to ClustalW classification). Nine α helices identified from the representative structure are marked at the bottom. Residues L60, S61, C65, R76, C77, H78 and V117 were marked with red squares.

The alignment of GLuc's homologous repeats [3,4]: residues 27-97 against residues 98-168 is shown in (B). Residues W143, L144, F151 [1] are marked with red circles, and the residues in the activity-associated loop R76-T79 [7] are marked with a red square. The flexible C-terminus is underlined. Clustal Co shows amino acid consensus generated by ClustalW (the symbol codes are the same as in (a)).
